# The replication checkpoint couples fork progression to the release of transcribed genes from nuclear pores

**DOI:** 10.64898/2026.10.07.757421

**Authors:** Giovanni Nigro, Jonathan Heuzé, Salomé Grassin, Lucie Fasolo, Antoine Barthe, Philippe Pasero, Jérôme Poli

## Abstract

The nuclear envelope contributes to chromatin compartmentalization in eukaryotes by anchoring heterochromatin to the lamina and active transcription sites to nuclear pore complexes (NPCs). During DNA replication, chromatin association with the nuclear envelope is thought to act as a topological barrier to DNA polymerases, potentially exacerbating transcription-replication conflicts (TRCs). While limited evidence suggests that the DNA replication checkpoint kinase Rad53^CHK1-2^ promotes the release of genes from the NPC during replicative stress, the global scale and precise regulation of this process remain unknown. Through a candidate-based screen, we demonstrate that gene release from NPCs is a general feature of the cellular response to replication stress, tightly coordinated with replication fork progression. We show that the checkpoint kinases Mec1^ATR^ and Rad53^CHK1^ are both essential for managing gene association with NPCs, but operate in a distinct, stepwise manner: Mec1^ATR^ triggers the initial dissociation by promoting the removal of RNAPII from the DNA, while Rad53^CHK1^ prevents the premature reassociation of transcribed loci by phosphorylating the nucleoporin Mlp1. Finally, we demonstrate that maintaining this control over genes-NPC association is critical for efficient DNA replication under stress. Collectively, our findings reveal that the DNA replication checkpoint integrates genome duplication with chromosome spatial positioning to maintain genome integrity during replication stress.

**Highlights:**

- Gene release from nuclear pores is coupled to replication fork progression
- Mec1-dependent RNAPII removal from chromatin triggers initial gene-NPC dissociation
- Rad53 phosphorylates Mlp1 to prevent premature gene reassociation with the NPC
- Dynamic control of gene-NPC contacts promotes efficient DNA replication under stress

## Introduction

During S phase, cells must duplicate their entire genome quickly and accurately. Throughout this process, replication forks frequently encounter various obstacles that impede their progression. This phenomenon is broadly termed replication stress, which is a hallmark of cancer cells (Saxena & Zou, 2022). Because transcription and replication occur on the same DNA strand, the transcriptional machinery represents one of the most prevalent endogenous obstacles to replication fork movement (Claussin et al., 2022; Gómez-González & Aguilera, 2019). Indeed, the loss of coordination between the transcription and replication programs is a major driver of oncogene-induced replicative stress and genome instability during the early stages of cancer onset (Macheret & Halazonetis, 2018). In response to replication stress, one of the main pathways to activate is the DNA replication checkpoint (DRC). This signaling cascade is initiated by the apical kinase Mec1^ATR^, which transduces the signal via the adaptor Mrc1^CLASPIN^ to the effector kinase Rad53^CHK1^. The DRC counteracts the adverse effects of replication stress through a concerted response including cell-cycle arrest, the suppression of late-origin firing, an increase in dNTP biosynthesis, and the stabilization and restart of stalled replication forks (Saxena & Zou, 2022). Although Rad53^CHK1^ is often considered the primary effector, Mec1^ATR^ also directly phosphorylates a distinct set of targets (Bastos de Oliveira et al., 2015; Hustedt et al., 2015), and both kinases perform critical, non-overlapping functions in maintaining chromosomal integrity.

Transcription acts as a pervasive source of obstacles for replication forks, manifesting as RNA:DNA hybrids (Hamperl et al., 2017; Heuzé et al., 2023), chromatin-bound RNA polymerases (Hurst et al., 2021; Poli et al., 2016a) and the accumulation of DNA supercoiling (Keszthelyi et al., 2016). The spatial organization of the genome can further exacerbate transcription-replication conflicts (TRCs) by preventing the dissipation of topological constraints. The nuclear envelope is a primary determinant of chromatin compartmentalization. While silent heterochromatin is often sequestered at the nuclear rim, transcribed euchromatin frequently contacts nuclear pore complexes (NPCs) – the specialized gateways for nucleocytoplasmic exchanges (Simon et al., 2024). Unlike the static association of heterochromatin to lamins, the spatial positioning of euchromatin is highly dynamic and adapts to the transcriptional landscape (Misteli, 2020). In budding yeast, numerous genes undergo a profound relocalization to physically associate with NPCs upon transcriptional activation (Brickner & Walter, 2004; Cabal et al., 2006; Casolari et al., 2004; Forey et al., 2021). Gene association with NPCs is mediated by two mechanisms: the nascent RNA recruits THO-TREX and the mRNA export complex Mex67 that interacts with nucleoporins (Nup1, Nup60, Mlp1/2, Dieppois, 2006) and TREX-2 that bridge RNAPII with nucleoporins (Nup1, Nup60) through interactions with SAGA (Cabal, 2006) and Mediator (Schneider, 2015). Similar movements have been observed in *C. elegans* for stress-response genes (Rohner et al., 2013), and in metazoans, where the association of enhancers with NPCs is required for cell-type-specific gene expression (Ibarra et al., 2016) and nucleoporins can directly contact genes in the nucleoplasm by dissociating from NPCs (Capelson et al., 2010; Kalverda et al., 2010). Although gene association to NPCs is not essential for transcription *per se*, it optimizes the kinetics of gene activation-repression (Ahmed et al., 2010; Schmid et al., 2006), promotes efficient mRNA processing (Gaillard et al., 2017; Kosar et al., 2021), and facilitates the inheritance of epigenetic information (Light et al., 2010). Beyond transcription, the NPC serves as a repair hub, for instance, persistent double-strand breaks (DSBs) relocalize to NPCs to channel specialized repair pathways in yeast and Drosophila (Horigome et al., 2014, 2016; Nagai et al., 2008; Ryu et al., 2015). Importantly, when mRNA processing is impaired in human cells, actively transcribed genes relocate to the nuclear periphery, where their persistent tethering triggers replication stress (Teloni et al., 2019). Thus, cells face a spatial conflict: the architecture that optimizes gene expression during interphase becomes a structural obstacle during S phase.

This constraint arises because DNA tethering to NPCs acts as a significant topological barrier. By anchoring the DNA template, the NPC restricts the free rotation required to dissipate transcription-and replication-induced supercoiling (Bermejo et al., 2012). This leads to an accumulation of hazardous topological constraints that would exceed the resolution capacity of topoisomerases (Keszthelyi et al., 2016), creating a roadblock for progressing replication forks. Despite its physiological importance, little is known about the mechanisms controlling DNA tethering to the nuclear envelope during DNA replication. In mouse embryonic fibroblasts, DNA-lamina interactions are disrupted during DNA replication (Lovejoy et al., 2023). In budding yeast, the association of inducible genes to NPCs is regulated by the Cdk1-mediated phosphorylation of the nucleoporin Nup1 (Garvey Brickner & Brickner, 2010).

Under conditions of replication stress, the DRC emerges as a central regulator of these spatial dynamics. Specifically, the effector kinase Rad53^CHK1^ directly phosphorylates the nucleoporin Mlp1^TPR^ to trigger the release of the *GAL1* locus from the NPC (Bermejo et al., 2011), while the apical kinase Mec1^ATR^ has been proposed to promote dissociation by inducing a transient shutdown of RNAPII transcription (Hurst et al., 2021). While these findings suggest that gene-NPC tethering is tightly regulated through the concerted action of multiple kinases, experimental evidence for gene release remains limited to a few inducible model genes like *GAL1*. This contrasts sharply with genome-wide studies indicating that over 600 constitutively expressed genes associate with NPCs during exponential growth (Forey et al., 2021), each representing a potential site of transcription-replication conflicts. Consequently, it remains unclear whether gene release from NPCs is a global response and how Mec1^ATR^ and Rad53^CHK1^ partition their roles to coordinate DNA spatial organization with the maintenance of replication integrity.

In this study, we conducted a targeted screen to characterize the spatial dynamics of constitutively expressed genes with respect to NPCs. We show that these highly expressed loci are systematically released from NPCs in response to hydroxyurea (HU)-induced replication stress in a checkpoint-dependent manner. By analyzing genes located at varying distances from active replication origins, we demonstrate that this release is not a global S-phase event, but is instead tightly coupled to replication fork progression. Paradoxically, we find that while constitutively expressed genes are released from NPCs during replication, HU-induced genes remain able to attach to NPCs, despite active checkpoint signaling. Furthermore, we reveal that the two main DRC kinases, Mec1 and Rad53, orchestrate this spatial organization through a distinct, two-step mechanism: Mec1^ATR^ triggers initial release by promoting the eviction of RNAPII, whereas Rad53^CHK1^ prevents premature reassociation by phosphorylating the nucleoporin Mlp1. Finally, we show that the failure to release these genes from NPCs impairs replication fork progression and sensitizes cells to replication stress. Our findings support a model in which the DRC dynamically reconfigures the spatial organization of the genome to facilitate the passage of replication forks through highly transcribed regions.

## Results

### Active genes are released from nuclear pore complexes during HU-induced replication stress

Given the fact that 10% of the genes associate with NPCs in budding yeast (Forey et al., 2021) and the established role of the DRC in releasing the inducible *GAL1-10* gene during replication stress (Bermejo et al., 2011), we sought to determine if gene-NPC dissociation is a general response to replication stress. To elucidate if other genes respond similarly to HU, we performed a candidate screen using strains that carry *lacO* repeats at various genomic positions. First, we analyzed the localization of candidate genes relative to the nuclear envelope in exponentially growing cells using the zoning assay (Fig. 1A). Among the loci tested, *ALG14* and *LSR1* showed no preferential enrichment at the nuclear periphery (Fig. 1B). While the internal position of *ALG14* is consistent with its low expression, *LSR1* remains non-peripheral despite being highly transcribed. This finding underscores that while high expression often correlates with NPC tethering, it is not by itself sufficient to drive NPC anchoring. In contrast, four other candidates -*RPL25*, *RPL29*, *RPL9A* and *TEF2* -exhibited robust association with the nuclear periphery, with 60-70% of cells displaying zone 1 enrichment (Fig. 1B). These results align with their status among the top 150 most highly expressed coding genes and their positive enrichment for Mlp1 by ChIP-seq (Forey et al., 2021). Next, we asked whether these constitutively tethered loci are released from NPCs in response to hydroxyurea (HU)-induced replication stress. Strikingly, 1-hour treatment with 0.2 M HU resulted in a strong reduction in the peripheral enrichment of all four highly expressed genes (*RPL25*, *RPL29*, *RPL9A* and *TEF2*). In these conditions, zone 1 occupancy dropped to ∼ 33%, a value consistent with a random spatial distribution (Fig. 1B). In contrast, HU treatment did not affect the localization of the non-peripheral control loci (*ALG14*, *LSR1*), which remained randomly distributed (Fig. 1B). These results suggest that the stress-induced release from the nuclear periphery is a widespread response of highly expressed genes.

**Figure 1:**
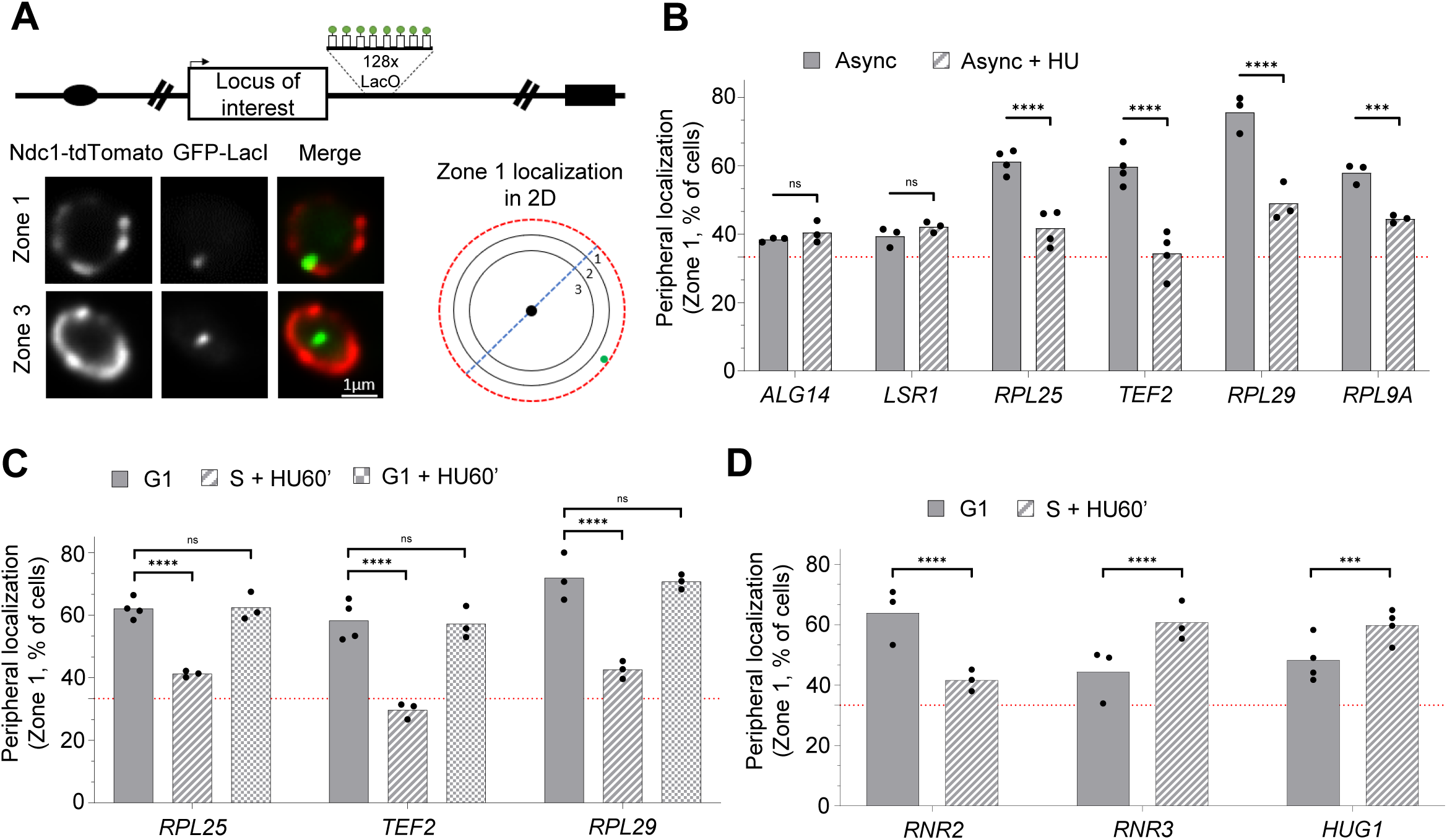
Actively transcribed genes are released from nuclear pores under HU-induced replication stress. (A) Schematic representation of a *lacO* array inserted in proximity to a gene of interest, which allows visualization via GFP-LacI. The nuclear periphery is visualized with Ndc1-tdTomato. Locus position is scored relative to the nuclear diameter in its plane of focus. Distance-over-diameter ratios are binned into three equal zones. 33% corresponds to a random distribution. Representative images show loci either positioned at the nuclear periphery (Zone 1) or in the nucleoplasm (Zone 3). (B-D). Percentage of GFP foci in Zone 1 under the indicated conditions. (B) Exponentially growing cells treated with 0.2 M hydroxyurea for 60 min (Async + HU) or left untreated (Async). (C-D) G1-arrested cells (G1), then cells released into S phase in the presence of 0.2M HU for 60min (S + HU60’) or kept in G1 arrest and treated with 0.2M HU for 60min (G1 + HU). For all panels (B-D), independent biological replicates are indicated by data points (N≥3, >100 cells/biological replicate). The horizontal dashed red line indicates 33% (random distribution). Statistical comparisons were performed using pairwise Fisher’s exact test (*P < .05, **P < .01, ***P < .001, ****P < .0001).

Exponentially growing cells include cells that are at different stages of the cell cycle, and HU is a potent inhibitor of dNTP biosynthesis that slows down DNA replication by ∼ 10-fold at 0.2 M (Poli et al., 2012). To confirm that HU treatment induces the release of transcribed genes by slowing down DNA replication, we first conducted the same experiment on a synchronized population by arresting cells in G1 with α-factor and releasing them into S phase + HU 0.2 M for 60 min. This analysis revealed no differences in the percentage of cells exhibiting a peripheral localization between exponentially growing cells and α-factor arrested G1 cells for *RPL25*, *RPL29* and *TEF2* (Fig. S1A). Releasing cells into S phase + HU 0.2M fully recapitulated the dissociation of genes from the nuclear periphery observed in non-synchronized cells treated with HU (Fig. 1C). To rule out an alternative effect of HU, we treated cells blocked in G1 with HU. As expected, HU in the absence of DNA replication did not release genes from the nuclear periphery (Fig. 1C). Consistently, treating G1 cells with HU did not activate the DNA replication checkpoint, contrary to S phase cells, as observed by monitoring changes in Rad53 mobility by Western blot (Fig. S1B). Although HU-induced replication stress shuts down housekeeping gene transcription, several genes are activated in response to HU treatment. To test if transcriptional activation is sufficient to promote the anchoring of genes to NPCs, we examined the position of *RNR2*, *RNR3* and *HUG1* in cells synchronized in G1 and released into S + HU for 60 min. We found that *RNR2* exhibits a strong peripheral localization in G1 cells (60%, Fig. 1D), whereas *RNR3* and *HUG1* are much lower (∼ 40%, Fig. 1D). This is consistent with the fact that *RNR3* and *HUG1* are only expressed upon replication stress, whereas *RNR2* is highly expressed even in the absence of HU (Elledge & Davis, 1990). In S phase cells submitted to HU, *RNR2* detaches from NPCs (∼ 40%, Fig. 1D); however, *RNR3* and *HUG1* increase their peripheral positioning (∼ 60%, Fig. 1D). We observed the same behaviour when exponentially growing cells are treated with HU (Fig. S1C). Taken together, these results argue that the release of highly expressed genes from NPCs is a general response to HU-induced replication stress that requires DNA replication itself to occur. However, this global dissociation does not prevent HU-inducible genes from being targeted to NPCs upon transcriptional activation.

### Gene release from NPC is coupled to replication fork progression

In yeast, the initiation of DNA replication follows a spatiotemporal program including ∼ 200 active and early origins and ∼ 250 origins that are activated later during S phase (Poli et al., 2012; Yabuki et al., 2002). Since the release of active genes from NPCs requires ongoing DNA replication, we reasoned that the dissociation could either occur all along the genome as soon as cells start to duplicate their genome or progressively with the firing of replication origins. To discriminate between these two possibilities, we inserted *lacO* repeats near genes at increasing distances from active early origins of replication. To identify loci that are associated with NPCs, we selected genes highly enriched in RNAPII (Rpb1 ChIP-seq, Forey et al., 2021) and found in proximity to NPCs (Mlp1 ChIP-seq; Forey et al., 2021). To preserve regulatory sequences, *lacO* repeats were inserted in an intergenic region proximal to the gene of interest (Fig. S2A). With this strategy, we generated integrations at 9 additional loci ranging from 1 to 120 kb from an active origin of replication (Fig. 2A). First, we analyzed the position of these loci with respect to the nuclear periphery using the zoning assay in exponentially growing cells. Consistent with their proximity to the NPC subunit Mlp1, all the loci tested showed a strong peripheral enrichment in microscopy (∼ 50 to 70 % peripheral, Fig. S2B). Arresting cells in G1 with α-factor did not significantly alter the peripheral enrichment regardless of the locus tested (∼ 60 to 70 % peripheral, Fig. 2B). Then, we scored the localization of the same loci when cells were released into S phase + 0.2 M HU for 60 min. Our analysis demonstrates that HU-induced replication stress triggers a distance-dependent release of genes from NPCs. Genes within 50 kb of an active origin show reduced peripheral localization (∼ 30 to 40 % peripheral, *PDC1*, *PYK2*, *AGP1*, *VMA13*, *TEF2*, *RPL29*, *SED1*, *RPL25*, and *PYK1*, Fig. 2B), whereas genes located more than 50kb remain largely attached to the NPC (∼ 60 to 70 % peripheral, *ACT1*, *ENO1*, *ZWF1*, Fig. 2B). Given that replication forks progress only 3-4 Kb from the origin at this dose of HU (Poli et al., 2012), our findings indicate that the release of genes from NPCs occurs prior to DNA duplication. This observation mirrors the fact that RNAPII is evicted from chromatin at distances reaching ∼30 kb from replication forks (Poli et al., 2016a; Tsirkas et al., 2022).

**Figure 2:**
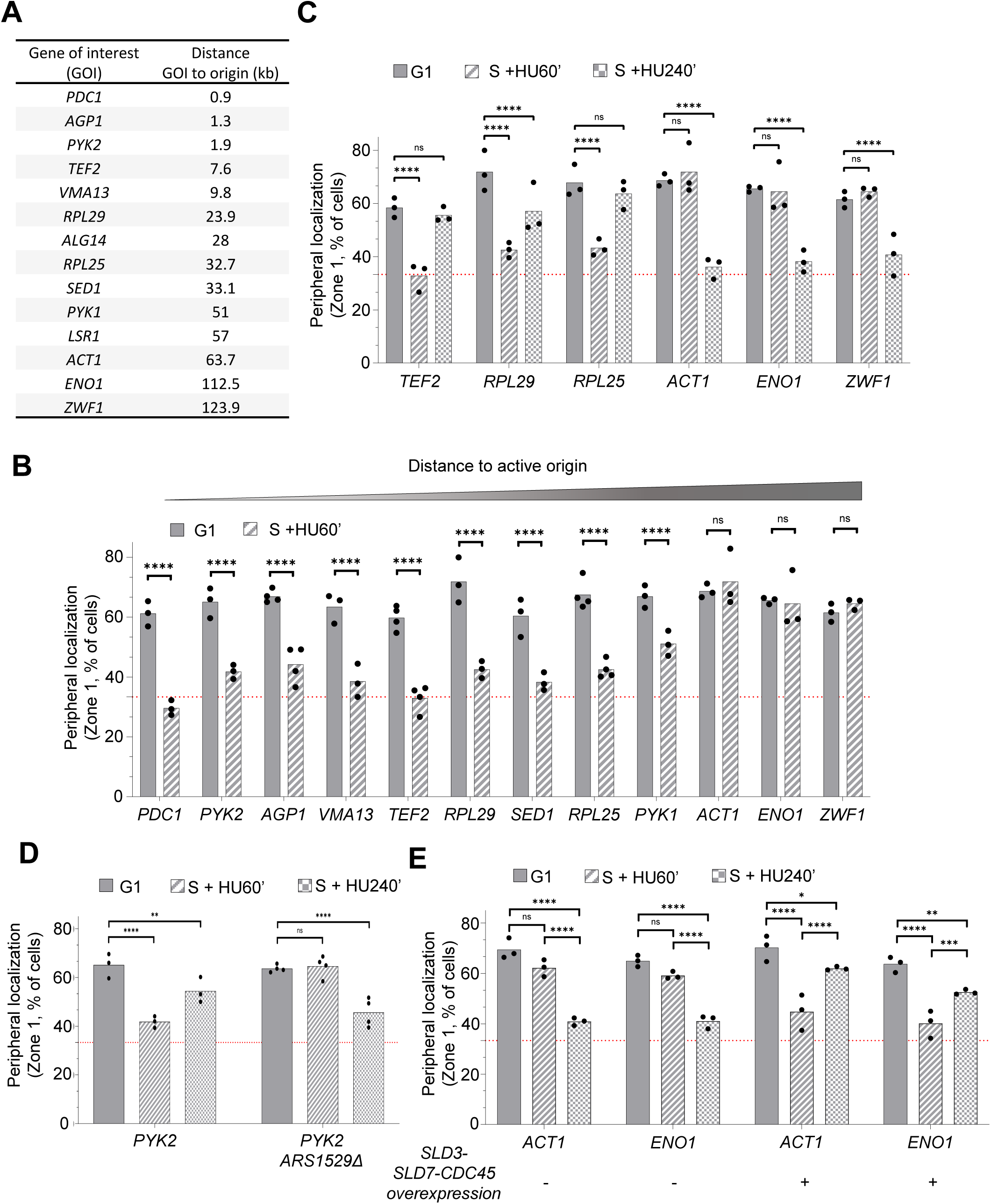
Gene release from nuclear pores is coupled to DNA replication progression. (A) Genes of interest targeted for *lacO* insertion, ranked by their distance to the closest active replication origin (in kb). (B) Percentage of GFP foci in Zone 1 for the indicated gene of interest, sorted by increasing distance from the closest active origin of replication in G1-arrested cells (G1) and then cells released into S phase in the presence of 0.2 M HU for 60 min (S + HU60’). (C-E) Percentage of GFP foci in zone 1 in G1-arrested cells (G1) then cells released into S phase in the presence of 0.2M HU for 60min (S + HU60’) or 240 min (S + HU 240’). (C) Wild-type (WT) cells. (D) Percentage of cells with the *PYK2* locus in zone 1 in WT or ARS1529-deleted (ARS1529Δ) cells. For (C) and (D), data for G1 and S + HU60’ are identical to those used in (B). (E) Strains carrying an estradiol-inducible overexpression of replication initiation factors (Sld3-Sld7-Cdc45). Overexpression is indicated by a (+). For all panels (B-D) Independent biological replicates are indicated by data points (N≥3, >100 cells/biological replicate). The horizontal dashed red line indicates 33% (random distribution).Statistical comparisons were performed using pairwise Fisher’s exact test (*P < .05, **P < .01, ***P < .001, ****P < .0001)

Our data suggest that the dissociation of genes from NPCs follows the spatiotemporal program of DNA replication. Given that replication forks are not stalled but are heavily slowed down under HU treatment, we hypothesized that genes scheduled for later replication would be released as the fork approaches or when late origins are activated. We addressed this possibility by tracking gene localization 240 minutes after releasing cells into S phase + 0.2 M HU. By this time point, replication forks typically progress ∼ 25 kb from each origin and later-firing origins will have fired (Poli et al., 2012). Consistent with our hypothesis, distal genes (>50 kb from early origins) were released from the nuclear periphery after 240 minutes into S phase + HU 0.2M (∼ 30 to 40 % peripheral, *ACT1*, *ENO1*, and *ZWF1*, Fig. 2C). Conversely, genes proximal to early origins (<50 kb) reassociated with NPCs, reaching ∼ 60 % peripheral localization (*TEF2*, *RPL29*, and *RPL25*, Fig. 2C). These observations suggest that gene positioning is a dynamic process during S phase, coordinated with the progression of DNA replication.

We further tested our model by manipulating the *PYK2* locus, which is positioned 3 kb from the active early origin ARS1529 in a region otherwise devoid of active origins (Fig. S2D). Deleting ARS1529 shifted *PYK2* to a late-replicating timing, leaving the locus over 50 kb away from the nearest active origin. In this experimental setup, *PYK2* failed to release from the NPC after 60 minutes of replication in 0.2 M HU but was instead detached from NPC after 240 minutes of HU treatment (Fig. 2D). On the other hand, in its natural context, *PYK2* reassociates with NPCs after 240 minutes of HU treatment, a behavior similar to the other origin-proximal genes *TEF2*, *RPL29*, and *RPL25* (Fig. 2C). Finally, we analyzed gene positioning in wild-type cells overexpressing limiting initiation factors (SLD3, SLD7 and CDC45) required for origin firing (Mantiero et al., 2011; Tanaka et al., 2011). In this context, the spatiotemporal program of origin firing is disrupted, causing late origins to fire prematurely in S phase. We overexpressed the initiation factors in strains carrying *lacO* arrays close to *ACT1* and *ENO1,* two genes that have late-firing origins at less than 5 kb (Fig. S2A). Remarkably, while *ACT1* and *ENO1* remained associated with NPCs in control cells after 60 minutes in HU, they were successfully released in cells overexpressing the initiation factors (Fig. 2E). Conversely, after 240 minutes into S phase + HU, the pattern reversed; cells overexpressing initiation factors displayed a reassociation of *ACT1* and *ENO1* with the NPC, whereas both loci were released from NPCs in control cells (Fig. 2E). We conclude that the release of transcribed genes from NPCs is a dynamic and reversible process. This release is triggered by advancing replication forks and propagates to loci up to ∼ 50 kb away. Furthermore, early-replicating genes can reassociate with NPCs later in S phase, even when genome duplication remains incomplete.

### Dynamic gene positioning allows efficient DNA replication

The observation that dozens – and potentially hundreds – of constitutively highly expressed genes undergo a coordinated release from NPCs suggests that this spatial reorganization is a critical response to replication stress. If gene tethering to NPCs acts as a physical barrier to the replication machinery, then preventing their release from the NPCs should sensitize cells to replication stress. We tested this hypothesis using two nucleoporin mutants, *mlp1-S1710A* and *nup1-S161D,* which are defective in releasing *GAL1-10* from NPCs during DNA replication (Bermejo et al., 2011; Garvey Brickner & Brickner, 2010). Zoning assays confirmed that in these mutants, two highly expressed genes *TEF2* and *RPL25* are associated with NPCs at similar levels to wild-type cells in exponentially growing cells (Fig. 3A). Consistent with the *GAL1-10* phenotype, neither mutant was able to release *TEF2* or *RPL25* upon HU treatment (Fig. 3A). Despite this persistent association with NPCs, these strains exhibited no significant sensitivity to chronic exposure to HU-induced replication stress in drop assays (Fig. 3B). Nevertheless, we investigated whether the inability to release genes from NPCs could impact the kinetics of DNA replication using DNA combing. We found that both mutants synthesized shorter BrdU tracks compared with wild-type cells (20.0 kb and 19.7 kb for *mlp1-S1710A* and *nup1-S341D*, respectively, vs 24.9 kb for WT; Fig. 3C). These results argue that persistent tethering to NPCs impede fork progression in cells that are checkpoint proficient.

**Figure 3:**
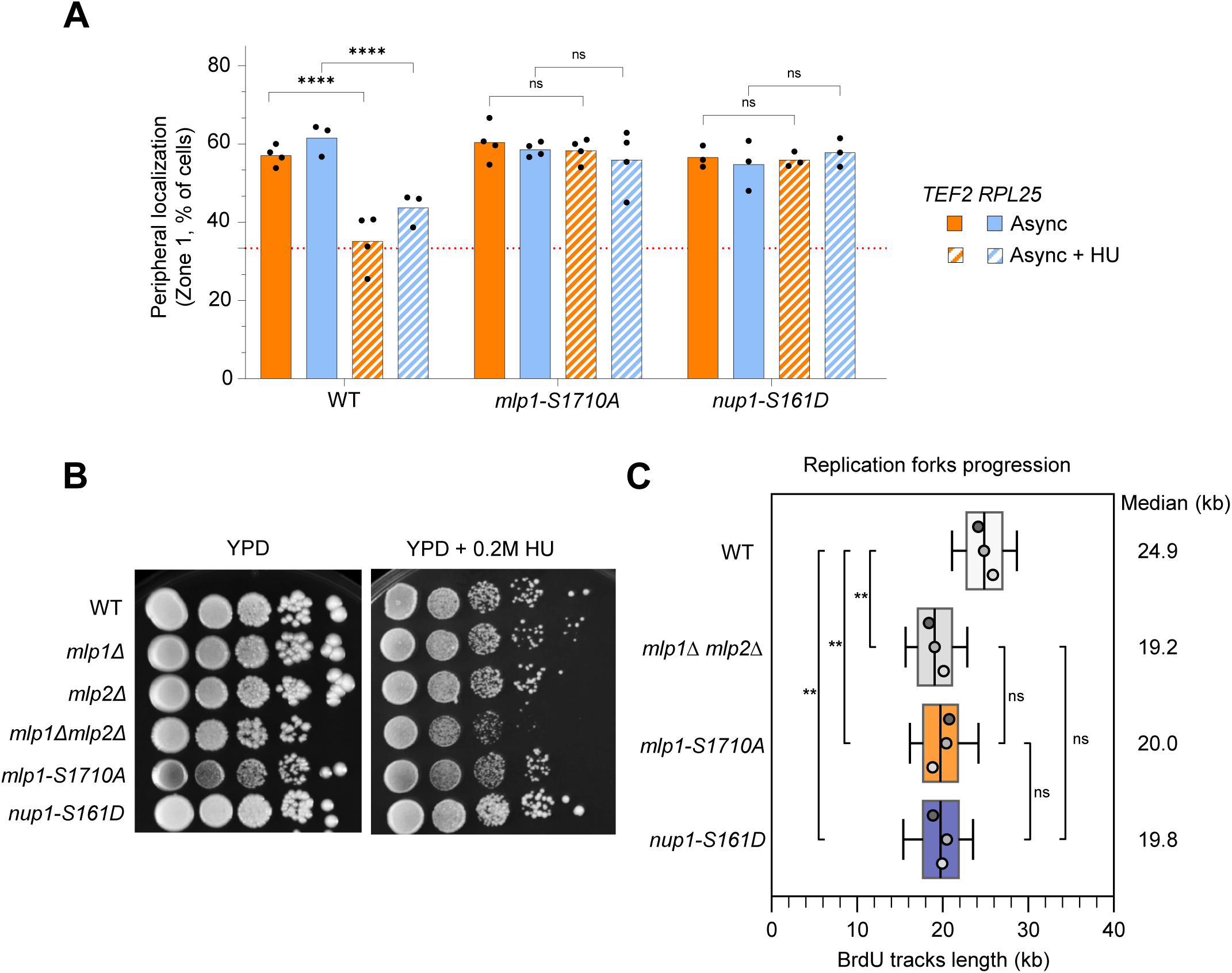
Dynamic control of genes association to nuclear pores promotes efficient DNA replication under HU stress. (A) Percentage of GFP foci in Zone 1 for *TEF2* and *RPL25* for the indicated strains in exponentially growing cells treated with 0.2 M hydroxyurea for 60 min (Async + HU) or left untreated (Async). Independent biological replicates are indicated by data points (N≥3, >100 cells/biological replicate). The horizontal dashed red line indicates 33% (random distribution). Statistical comparisons were performed using pairwise Fisher’s exact test (*P < .05 and ****P < .0001) (B) Ten-fold serial dilutions of cells of the indicated strains were spotted onto YPD +/- 0.2 M HU. Each assay was performed at least twice; one representative experiment is shown. (C) Analysis of replication fork progression at the single-molecule level by DNA combing. Exponentially growing cells were synchronized in G1 with α-factor and released into S phase in the presence of 0.2M HU. Newly replicated DNA was labeled with BrdU for 180 min, added 60 min after release into S phase + HU. The graph depicts the distribution of BrdU track length. Box, 25-75 percentile range. Whiskers, 10-90 percentile range. Median is indicated in kb. A one-way ANOVA was performed on replicate median values, followed by Tukey’s multiple-comparisons test. Significance is denoted as ns ≥ 0.05, *P < 0.05 and **P < 0.001. 3 biological replicates are shown.

In yeast, the vast majority of highly expressed genes associate with NPCs during exponential growth, a spatial organization that is abolished in the absence of the nucleoporins Mlp1 and Mlp2 (Forey et al., 2021). While our previous results highlighted the necessity of releasing constitutively expressed genes from NPCs during HU stress, we also observed that HU-induced genes actively relocate to NPCs during DNA replication (Fig. 2). This dual behaviour prompted us to examine the impact of abrogating all gene-NPC contacts upon HU exposure. To this end, we used the *mlp1Δmlp2Δ* double mutant, which completely loses gene-NPC contacts. While this mutant exhibited only minor sensitivity to chronic HU exposure in drop assays (Fig. 3B), it displayed a striking defect in replication fork progression. Specifically, DNA combing revealed a reduction in fork rate similar to that observed in *mlp1-S1710A* (19.2 vs 20.0 kb; Fig. 3C). Together, these findings suggest that loss of gene-NPC tethering correlates with impaired replication fork progression under replication stress. Our data thus point to a dynamic spatiotemporal program of gene-NPC association promotes replication under HU stress. This underscores that NPC tethering not only represents a topological constraint but also contributes to an efficient response to replication stress.

### Mec1 and Rad53 control gene-NPC contacts through independent mechanisms

We and others have shown that the DNA replication checkpoint (DRC) triggers the dissociation of *GAL1-10* genes from NPCs during HU-induced replication stress (Bermejo et al., 2011; Hurst et al., 2021). However, Mec1^ATR^ and Rad53^CHK1^ facilitate gene release through distinct mechanisms: Rad53^CHK1^ acts directly on the NPC by phosphorylating the nucleoporin Mlp1^TPR^ (Bermejo et al., 2011), whereas Mec1^ATR^ promotes release indirectly by stimulating the removal of RNAPII from chromatin (Hurst et al., 2021). The existence of these dual pathways raises the question of whether they function redundantly or if they have distinct roles. To understand the role of DRC kinases in the release of highly expressed genes from NPCs, we first compared the peripheral localization of *RPL25* and *TEF2* in wild-type cells and several DRC mutants, in the presence and absence of HU. This analysis revealed that all DRC mutants failed to release both genes in response to HU treatment (Fig. 4A). To test if Rad53-dependent phosphorylation of Mlp1 at S1710 is sufficient to release genes from NPCs, we scored *RPL25* and *TEF2* peripheral localization in DRC mutants expressing the phosphomimetic allele *mlp1-S1710D*. For that, we chose the Rad53 kinase-deficient mutant *rad53-11* (Weinert, 1994) and the Mec1 kinase-deficient mutant *mec1-100* that specifically impairs Rad53 activation during replication stress (Paciotti, 2006). We note that *mlp1-S1710D* is proficient to tether both genes at NPCs in the absence of replication stress, and for release during HU treatment (Fig. 4A). As previously reported, the *mlp1-S1710D* rescues the gene-release defect in *rad53-11* cells (Fig. 4A). We observed a similar rescue in *mec1-100* background, an allele that impairs Rad53 activation during replication stress and consequently prevents Mlp1-S1710 phosphorylation (Fig. 4A). Contrary to *mec1-100* and *rad53-11*, the non-phosphorylatable allele *mec1-S1991A* is fully proficient for activating Rad53 upon HU treatment but fails to evict RNAPII from chromatin (Hurst et al., 2021). Combining the *mlp1-S1710D* allele with the *mec1-S1991A* mutant failed to rescue the gene release defect in the *mec1-S1991A* background (Fig. 4A). This indicates that *mec1-S1991A* is epistatic to Mlp1 phosphorylation by Rad53^CHK1^, suggesting that RNAPII eviction from chromatin is essential for gene release from NPCs, regardless of Mlp1 phosphorylation status. To test if RNAPII retention on chromatin causes the failure to detach genes from NPCs in cells lacking a functional Mec1^ATR^, we used the anchor away system to force nuclear eviction of Rpb1, the large subunit of RNAPII. In this system, the eviction of Rpb1 is inducible by the addition of rapamycin (Haruki et al., 2008). Cells bearing the anchor-away background, including the deletion of *SML1* that is required for *mec1Δ* viability, readily anchor *RPL25* to the nuclear periphery in the absence of rapamycin (WT, *sml1Δ* and *mec1Δ sml1Δ* ∼ 60 %, Fig. 4B). Upon HU treatment, checkpoint-proficient cells released *RPL25* (WT and *sml1Δ* ∼ 30 %, Fig. 4B) whereas *mec1Δsml1Δ* mutant failed to release this gene (*mec1Δsml1Δ* ∼ 60 %, Fig. 4B). Forced Rpb1 eviction (+ rapamycin) released *RPL25* in exponentially growing cells and in *mec1Δsml1Δ* strains exposed to HU (∼ 30 %, Fig. 4B). These findings support the view that Mec1-dependent removal of RNAPII from chromatin is sufficient to release genes from NPCs.

**Figure 4:**
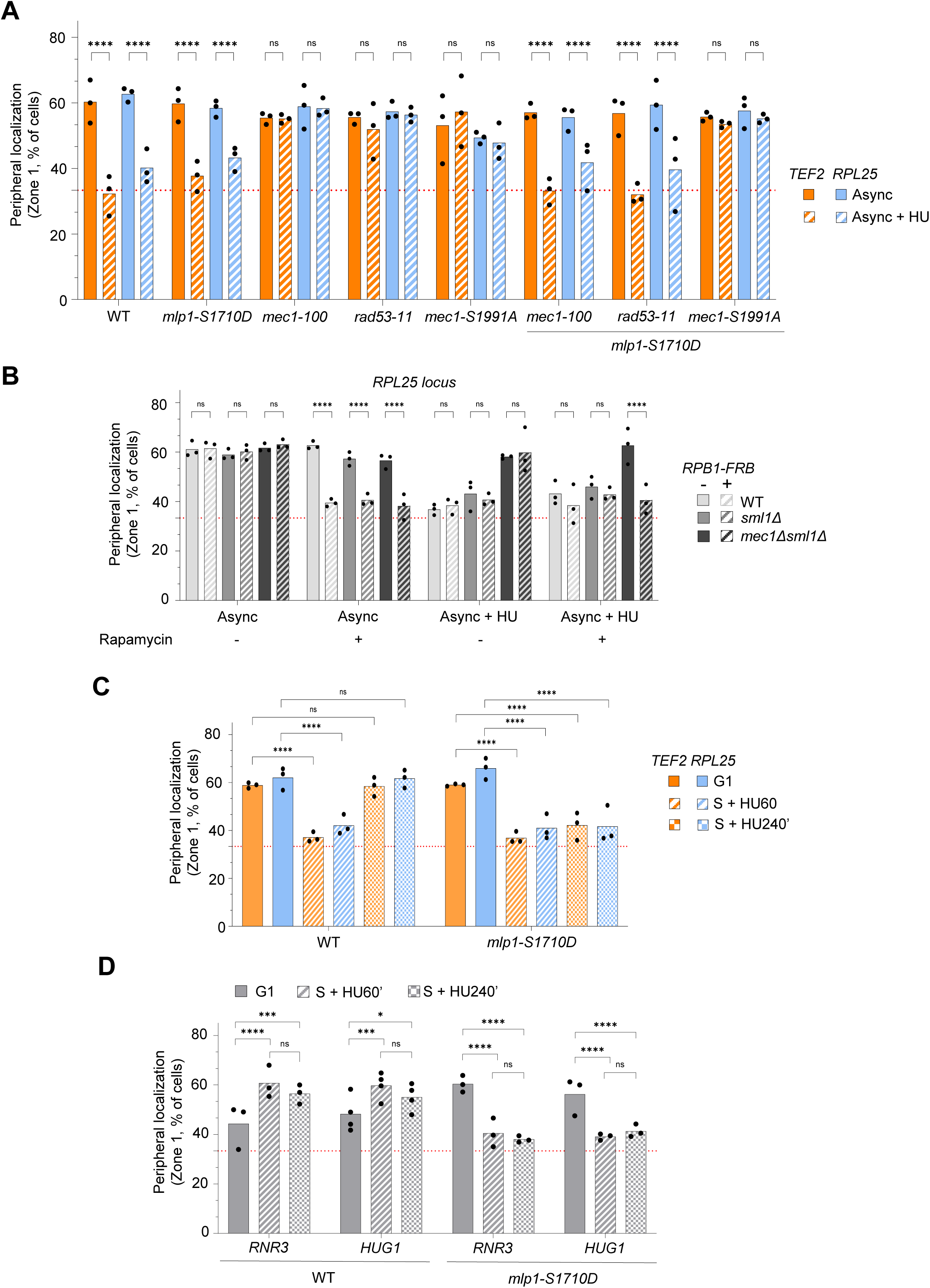
Mec1^ATR^ and Rad53^CHK1^ have independent roles in the release of genes from NPCs during replication stress. (A) Percentage of GFP foci in Zone 1 for *TEF2* and *RPL25* for the indicated strains in exponentially growing cells treated with 0.2 M hydroxyurea for 60 min (Async + HU) or left untreated (Async). (B) Rpb1 anchor-away strains in which the large subunit of RNAPII (Rpb1) can be evicted from the nucleus upon addition of rapamycin (1ug/ml). Percentage of GFP foci in zone 1 for *RPL25* for the indicated strains in exponentially growing cells treated with 0.2M hydroxyurea for 60 min (Async + HU) or not (Async) +/- rapamycin 30 min before collection. (C-D) Percentage of GFP foci in Zone 1 in G1-arrested cells (G1) and cells released into S phase in the presence of 0.2 M HU for 60 min (S + HU60’) or 240 min (S + HU 240’). *(C) TEF2* and *RPL25* or (D) *RNR3* and *HUG1* are shown in the indicated strains. For all panels (B-D) Independent biological replicates are indicated by data points (N≥3, >100 cells/biological replicate). The horizontal dashed red line indicates 33% (random distribution). Statistical comparisons were performed using pairwise Fisher’s exact test (*P < .05, **P < .01, ***P < .001, ****P < .0001).

Given that Mlp1-S1710 phosphorylation does not trigger gene release when RNAPII is still present on chromatin, we sought to determine if it has a distinct role during replication stress. Previously, we observed that early-replicated genes reassociate with NPCs after 4 h in S phase + HU (Fig. 2). Thus, we asked whether the phosphorylation of Mlp1 could control reassociation of genes to NPCs by quantifying the peripheral localization of *TEF2* and *RPL25* in *mlp1-S1710D* cells released into S phase + HU for 1 h and 4 h. While both wild-type and *mlp1-S1710D* strains successfully released *TEF2* and *RPL25* after 1 h in HU, the *mlp1-S1710D* mutant exhibited a striking failure to reassociate either locus with NPCs after 4 h in HU (Fig. 4C). This observation suggests that the *mlp1-S1710D* allele prevents reassociation of genes with NPCs while cells are still replicating. If this model is correct, we reasoned that *mlp1-S1710D* might also preclude NPCs association of HU-induced genes. We found that *RNR3* and *HUG1* associate with NPCs in wild-type cells treated with HU after release into S phase for 1 h and 4 h but remained detached under the same conditions in cells bearing the *mlp1-S1710D* allele (Fig. 4D). Similar results were obtained in exponentially growing cells treated with HU (Fig. S3A). Collectively, these results point toward distinct roles for Mec1^ATR^ and Rad53^CHK1^ in the control of gene association with NPCs during replication stress.

### RNAPII occupancy and gene-NPC tethering are additive sources of transcription-replication conflicts in checkpoint mutants

S-phase checkpoint mutants are acutely sensitive to acute and chronic exposure to HU-induced replication stress, a phenotype that stems partly from unresolved transcription-replication conflicts (TRCs). For *rad53* mutants, part of this sensitivity stems from the accumulation of topological constraints caused by persistent gene-NPC association (Bermejo et al., 2011). In contrast, *mec1* mutants accumulate RNAPII on chromatin, which directly hinders replication fork progression (Hurst et al., 2021; Poli et al., 2016a). Because gene tethering at NPCs requires active RNAPII (Dieppois & Stutz, 2010), these two sources of TRCs are mechanistically intertwined, making it difficult to distinguish between the impact of RNAPII as a roadblock and the topological barriers generated by NPC tethering. To investigate the respective impacts of RNAPII occupancy and NPC tethering, we undertook a genetic approach in *mec1* and *rad53* mutants. We used a C-terminal TAP-tagged allele of *RPB3* to reduce RNAPII occupancy on chromatin without impacting cell growth (Hurst et al., 2021), and *mlp1-S1710D* to bypass checkpoint signaling and force gene release from NPCs. First, we looked for genetic suppression upon chronic exposure to HU using drop assays. Growth of *RPB3-TAP* and *mlp1-S1710D* strains was indistinguishable from isogenic wild-type cells in the absence or presence of HU (Fig. 5A-C). We selected *mec1-S1991A*, as this mutant exhibits slow fork progression, fails to decrease the occupancy of RNAPII on chromatin and fails to detach genes tethered to NPCs. This mutant was readily rescued by the addition of *RPB3-TAP* but not by *mlp1-S1710D* (Fig. 5A). Conversely, the *mec1-100* mutant which impairs Rad53 activation in HU but retains some Mec1-specific functions was only rescued by *mlp1-S1710D* (Fig. 5B). The kinase-deficient *rad53-11* mutant was suppressed by *RPB3-TAP* and *mlp1-S1710D* (Fig. 5C). The combination of *RPB3-TAP* and *mlp1-S1710D* did not further improve the growth of any checkpoint mutant upon chronic exposure to HU (Fig. 5A-C). These genetic suppressions suggest that TRCs in checkpoint mutants emanate from RNAPII acting as a direct roadblock and from topological barriers mediated by association with NPCs.

**Figure 5:**
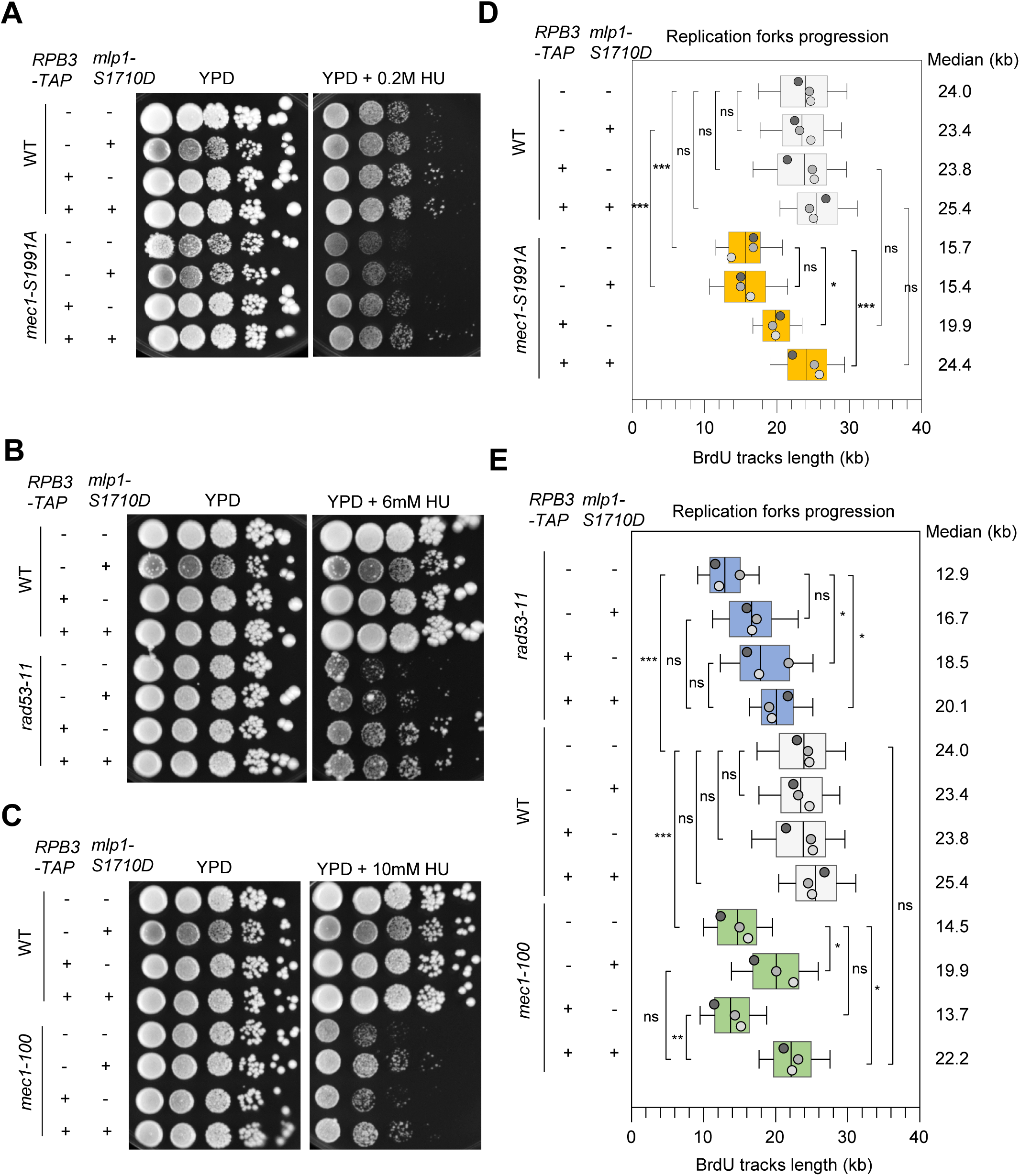
RNAPII occupancy and gene association to nuclear pores function as additive sources of transcription-replication conflicts in checkpoint mutants. (A, B, C) Ten-fold serial dilutions of cells of the indicated strains were spotted onto YPD +/- the indicated HU doses. Each assay was performed at least twice; one representative experiment is shown. (D, E) Analysis of replication fork progression at the single-molecule level by DNA combing. Exponentially growing cells were synchronized in G1 with α-factor and released into S phase in the presence of 0.2 M HU. Newly replicated DNA was labeled with BrdU for 180 min, added 60 min after release into S phase + HU. The graph depicts the distribution of BrdU track length. Box, 25-75 percentile range. Whiskers, 10-90th percentiles range. Median is indicated in kb. A one-way ANOVA performed on replicate median values followed by Tukey’s multiple-comparisons test. Significance is denoted as ns ≥ 0.05, *P < 0.05, **P < 0.001 and ***P < 0.0001 3 biological replicates are shown. Data for WT, RPB3-TAP, mlp1-S1710D and RPB3-TAP mlp1-S1710D are the same between Fig5D and fig5E

To see if the reduction of TRCs by either *RPB3-TAP* or *mlp1-S1710D* rescues the DNA replication defects of checkpoint mutants upon HU, we performed DNA combing. Consistent with earlier results showing the suppression of replication fork progression defects, we found that *RPB3-TAP* partly suppresses the slow-fork phenotype observed in the *mec1-S1991A* mutant (19.9 vs 15.7 kb, Fig. 5D, Hurst, 2021) whereas the addition of *mlp1-S1710D* did not improve replication in this mutant (15.4 vs 15.7 kb, Fig. 5D). This result aligns with the fact that *mlp1-S1710D* did not rescue the release of genes from NPCs in *mec1-S1991A* (Fig. 4A). Remarkably, the simultaneous introduction of *RPB3-TAP* and *mlp1-S1710D* into *mec1-S1991A* restored fork progression to wild-type levels (24,8 vs 24.4 kb, respectively, Fig. 5D). This suggests an additive suppression of the slow-fork phenotype under HU stress in which decreasing RNAPII occupancy and preventing reassociation of genes with NPCs alleviates replication stress.

Next, we sought to determine how the reduction of RNAPII occupancy (*RPB3-TAP*) synergizes with the relief of NPC-mediated topological constraints (*mlp1-S1710D*) to rescue replication stress in checkpoint mutants that are more sensitive to HU. To this end, we compared replication fork progression in *mec1-100* and *rad53-11* expressing one or both suppressors. The introduction of either allele partially rescued slow fork progression in *rad53-11* (Fig. 5F) whereas only *mlp1-S1710D* improved fork progression in *mec1-100* (Fig. 5E). Then, we examined each mutant combined with both suppressors and found a slight additive improvement in replication fork progression for *rad53-11* and *mec1-100* (20.1 vs 18.5 kb for *rad53-11* and 22.2 vs 19.9 kb for *mec1-100*, Fig. 5E). Collectively, these findings support the view that transcription-replication conflicts in checkpoint mutants arise from the combined effects of RNAPII occupancy and gene-NPC association.

## Discussion

Transcription is the major endogenous impediment to replication fork progression, and the DNA replication checkpoint is essential for minimizing the deleterious effects of transcription-replication conflicts (Landsverk et al., 2019; Poli et al., 2016b). In *S. cerevisiae*, the checkpoint kinases Mec1^ATR^ and Rad53^CHK1^ orchestrate a multi-layered response through distinct yet complementary pathways. While Mec1^ATR^ induces the eviction of RNAPII from chromatin to clear the path for DNA synthesis (Hurst et al., 2021; Poli et al., 2016b), Rad53^CHK1^ has been implicated in the release of transcribed genes from nuclear pore complexes (Bermejo et al., 2011). However, this latter function was previously characterized only for the inducible *GAL1-10* locus, leaving it unclear whether NPC dissociation is a general response to replication stress and to what extent this anchoring contributes to the severity of TRCs.

Our results demonstrate that the dissociation of actively transcribed genes from NPCs is a global response to replication stress, under the control of the DNA replication checkpoint (Fig. 6). Crucially, this release is not triggered simultaneously across the genome, rather it is spatially and temporally coupled to the progression of replication forks. This suggests that the checkpoint does not simply clear gene-NPC tethering at the onset of S phase, but instead fine-tunes this spatial organization as replication progresses. Paradoxically, we find that HU-induced genes retain the ability to associate with the NPCs even under strong checkpoint activation, revealing a selective control mechanism that balances the need for stress-responsive transcription with the maintenance of replication integrity. We propose that checkpoint-dependent gene release from NPCs follows a two-step mechanism. Our data support a hierarchy in which Mec1^ATR^-dependent RNAPII eviction is the initiating event. In this model, RNA polymerase removal is necessary to release the transcribed locus from the NPC. During HU-induced replication stress, degradation of the RNAPII large subunit Rpb1 on chromatin (Poli, 2016, Hurst, 2021) likely releases transcribed genes from NPCs by disrupting both RNA-dependent (THO/TREX/Mex67) and protein-mediated (TREX-2/Mediator/SAGA) interactions with nucleoporins. While Mec1-induced Rpb1 degradation requires INO80, the Paf1 complex, and the proteasome (Poli, 2016, Lafon, 2015), the precise mechanism connecting Mec1-dependent phosphorylation to RNAPII degradation remains to be elucidated. However, transcription is a stochastic and recurring process. Without a secondary safeguard, the transcribed locus could rapidly reassociate with NPCs. Rad53^CHK1^ fulfills this function by phosphorylating the nucleoporin Mlp1, which prevents the premature reassociation of transcribed loci. Our findings that replication defects in the *mec1-S1991A* mutant are additively rescued by both reducing RNAPII occupancy and disrupting NPC association provide key insights into the nature of TRCs. This leads us to a model in which RNAPII occupancy and NPC tethering function as distinct but additive sources of stress: the RNA polymerase likely acts as a direct physical roadblock, while gene-NPC anchoring imposes a topological constraint that further prevents its bypass or resolution. By coordinating both the eviction of the transcriptional machinery and the association with NPCs, the DNA replication checkpoint kinases resolve these convergent impediments to safeguard genome integrity.

**Figure 6:**
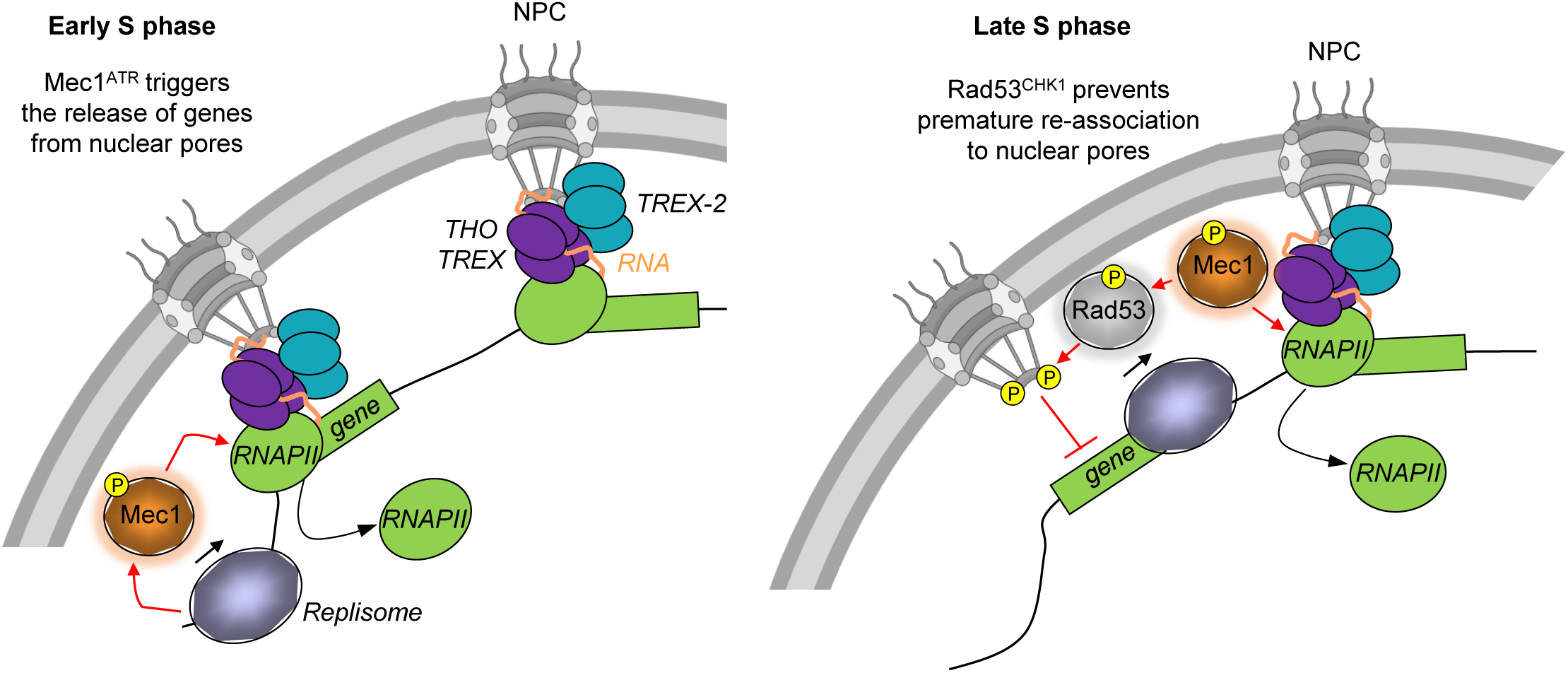
The DNA replication checkpoint kinases Mec1^ATR^ and Rad53^CHK1^ avoid transcription-replication conflicts by coupling replication fork progression to transcription inhibition during replication stress. In yeast, a majority of highly transcribed genes associate physically with nuclear pores, which represent topological barriers to ongoing replication forks. We propose that the DNA replication checkpoint promotes fork progression through two distinct mechanisms: (1) Mec1^ATR^ triggers the initial release of transcribed genes from nuclear pores by promoting the eviction of RNAPII from chromatin, and (2) Rad53^CHK1^ prevents the premature reassociation of transcribed genes to nuclear pores by directly phosphorylating Mlp1^TPR^. Both mechanisms contribute to the efficient replication of genomic DNA during HU stress. Finally, our data suggest that the control of gene-nuclear pore association by the checkpoint is coupled to replication fork progression in space and time.

We found that the release of genes from NPC is enhanced during HU-induced replication stress compared to unperturbed S phase, with only one of the three gene being released under physiological conditions. Although basal S-phase replication stress induces low-level activation of Mec1 and Rad53, exposure to HU triggers a potent checkpoint kinase signaling. This suggest that only a subset of genes is targeted for NPC-release during an unperturbed S phase or that release events occur with lower frequency in the absence of exogenous stress.

While our study focused on the Rad53^CHK1^-dependent targeting of Mlp1 at S1710, it is clear that gene-NPC association is governed by parallel mechanisms throughout the cell cycle. For instance, Cdk1-mediated phosphorylation of Nup1 at S161 promotes the tethering of inducible genes, such as *GAL1-10* and *INO1* during G1 and G2, while the loss of this phosphorylation during S phase facilitates their release (Garvey Brickner & Brickner, 2010). We found that the expression of either a phosphomimetic *nup1-S161D* allele or a non phosphorylatable *mlp1-S1710A* allele was sufficient to block the dissociation of highly constitutively expressed genes like *TEF2* and *RPL25*. The fact that a mutation in either Nup1 or Mlp1 leads to a similar failure to release these loci from NPCs suggests a dominant negative effect. This implies that these two regulatory events are not redundant but rather converge to allow the efficient release of genes from NPCs. Consistent with this hypothesis, both phosphomutants exhibit comparable replication defects under HU stress. The control of transcribed gene association with nuclear pores by CHK1^Rad53^ to facilitate replication fork progression under stress appears to be a highly conserved mechanism in eukaryotes (accompanying manuscript by Wen *et al*.).

Our study reveals that gene-NPC dissociation does not occur all at once upon exposure to replication stress. Instead, its dissociation is tightly coupled to the progression of replication forks across the genome. By analyzing loci at varying distances from active origins, we show that the release occurs progressively, extending up to ∼ 30 kb from active replication sites. This localized response mirrors the propagation of RNAPII eviction previously reported (Poli et al., 2016b; Tsirkas et al., 2022). Indeed, we demonstrate that forcing the eviction of Rpb1, the large subunit of RNAPII, is sufficient to disrupt the NPC association of the *RPL25* locus, even in *mec1* checkpoint mutants. This is consistent with the fact that active transcription machinery is the primary tether anchoring genes to NPCs (Dieppois & Stutz, 2010) and suggests RNAPII eviction and NPC detachment are functionally integrated events. Consistent with a replication-dependent mechanism, we observed that the addition of HU to cells arrested in G1 phase, where no replication forks are active, fails to trigger the release of these genes from the NPC. We propose that the signaling cascade originates at stalled forks, where ssDNA-RPA accumulation recruits and activates the Ddc2^ATRIP^-Mec1^ATR^ (Hustedt et al., 2013). While the precise mechanism of signal propagation remains to be elucidated, we hypothesize that the physical state of the DNA template plays a role. During HU-induced replication stress, dNTP shortage stalls DNA polymerases, while the MCM helicase continues to unwind the double helix. This functional uncoupling not only generates the ssDNA required for checkpoint activation but also creates a wave of positive supercoiling (Keszthelyi et al., 2024) that may act as a long-range mechanical signal triggering RNAPII eviction and NPC release ahead of the advancing replisome. From an evolutionary perspective, triggering the removal of potential transcription-replication conflicts before the fork physically encounters RNAPII – whether associated with NPCs or not – likely promotes the maintenance of genome stability.

Interestingly, we show that the release of active genes from NPCs is not definitive, but rather a transient state linked to the passage of replication forks. Even as the checkpoint remains active, we observe that certain loci, specifically the HU-responsive genes *RNR3* and *HUG1,* can establish NPC tethering during replication stress. Furthermore, early-replicating genes that dissociate at the onset of HU stress (S + HU 60 min) exhibit re-attachment to NPCs as cells progress further into S phase (S + HU 240 min). These findings suggest that the checkpoint-dependent clearance of RNAPII from the DNA template is localized and temporary. Notably, *RNR3* and *HUG1* are located 7 and 21 kb from an active origin, respectively, well within the ∼ 30 kb range in which constitutively expressed genes are released. This implies that the massive transcriptional induction of stress-response genes can override the general release mechanism. Intriguingly, we found that expressing the phosphomimetic allele *mlp1-S1710D* prevents this association throughout the HU treatment, suggesting that the presence of a constitutively phosphorylated Mlp1 locks the NPC in a non-permissive state. In wild-type cells, this implies that Rad53-mediated phosphorylation of Mlp1 is either temporally fine-tuned during replication stress or limited to a subset of nuclear pores. This latter possibility is supported by recent evidence of structural heterogeneity within the NPC population, with a distinct subset of nuclear pores being deprived of Mlp1 (Bensidoun et al., 2022). Crucially, these Mlp1-deficient NPCs have been shown to interact with Ctf18, a factor essential for PCNA loading and checkpoint signaling (Choudhry et al., 2023). Further experiments are required to determine whether stress-induced genes associate specifically with a subpopulation of NPCs.

Beyond its role as a topological barrier, association with the NPC constitutes a specialized environment for maintaining RNA:DNA hybrid homeostasis. Evidence from both yeast and human cells suggests that the association of actively transcribed genes with NPCs limit the accumulation of RNA:DNA hybrids (Gaillard et al., 2017; Kosar et al., 2021; Penzo et al., 2023). Additionally, loci prone to R-loop accumulation are actively repositioned to the NPC via an RPA-SUMO-dependent pathway, a recruitment mechanism similar to that employed by collapsed replication forks and persistent DNA lesions (Penzo et al., 2023; Schirmeisen et al., 2024). This recruitment likely facilitates the enzymatic dissociation of RNA:DNA hybrids. For instance, Ddx19, an RNA:DNA hybrid helicase, interacts with NPC subunits after an ATR^Mec1^-dependent nuclear translocation (Hodroj et al., 2017). Thus, under replication stress, the relocation of stress-induced genes to the NPC could constitute a mechanism for limiting transcriptional R-loops during DNA replication. Consistently, the DNA replication checkpoint sorts the genome to sustain replication fork progression while ensuring a proper replication stress response.

The proposal that RNAPII and NPC tethering act as additive barriers for replication forks is consistent with recent findings at the copper-resistance gene *CUP1* (Whale et al., 2022). Upon chronic exposure to high levels of copper, the transcriptional activation of *CUP1* triggers adaptive copy number amplification of the locus, thereby increasing resistance to copper. They found that while high levels of transcription at *CUP1* induce TRCs and stall replication forks, copy number variation occurs specifically in cells proficient in tethering the *CUP1* gene to NPCs. In this context, while high RNAPII occupancy induces fork stalling, association with the NPC channels these stalled forks into a Rad51-Rad52-dependent homologous recombination pathway. Intriguingly, unlike the constitutively expressed loci in our study, *CUP1* appears to escape the transcription control of the DNA replication checkpoint, as the loss of Mec1^ATR^ does not impact copy number variation. This implies that while persistent tethering to the NPC jeopardizes replication fork stability and genome duplication, it can also function as a potent driver of genome plasticity. Thus, gene-NPC tethering represents a double-edged sword: while it is a source of replication stress that must be counteracted by the checkpoint to maintain genome stability, it also promotes survival mechanisms through adaptive gene amplification.

## Supplemental Figure legends

**Figure S1:**
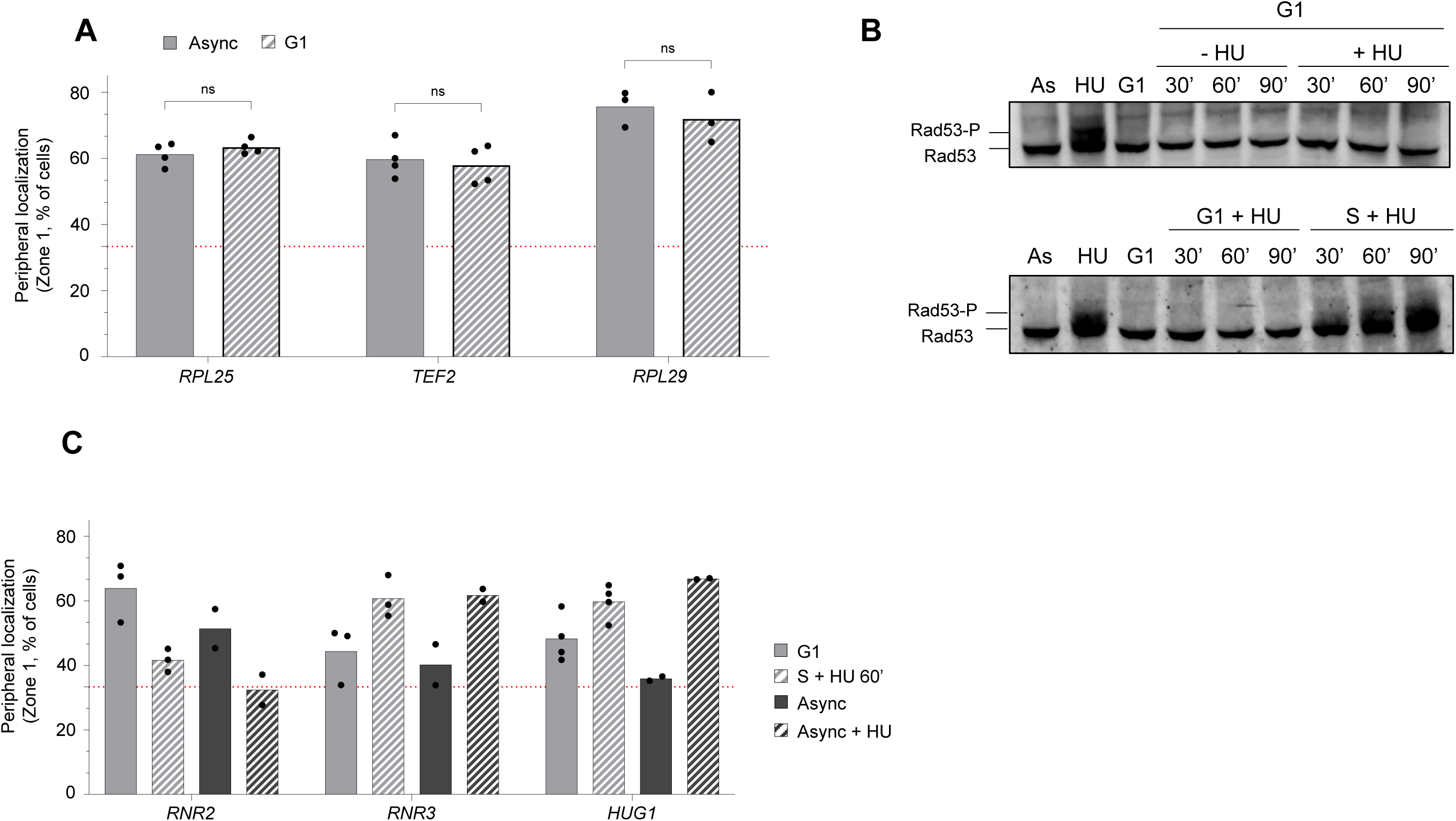
Actively transcribed genes are released from nuclear pores under HU-induced replication stress. (A and C) Percentage of GFP foci in Zone 1 under the indicated conditions (Async = exponentially growing cells, Async + HU = exponentially growing cells treated with 0.2 M HU for 60 min, G1 = arrested with α-factor, S+HU60 synchronously released cells in S phase treated with 0.2 M HU for 60 min). For all panels, independent biological replicates are indicated by data points (N≥2, >100 cells/biological replicate). The horizontal dashed red line indicates 33% (random distribution). Statistical comparisons were performed using a pairwise Fisher’s exact test only for experiments with n≥3 (*P < .05, **P < .01, ***P < .001, ****P < .0001). (B) Exponentially growing cells treated with 0.2 M HU (HU) or left untreated (AS). G1-arrested cells (G1) or cells treated with 0.2 M (+HU) or left untreated (-HU). Rad53 phosphorylation was detected by immunoblotting using a polyclonal anti-Rad53 antibody. The arrow indicates phosphorylated Rad53, detectable as a mobility shift.

**Figure S2:**
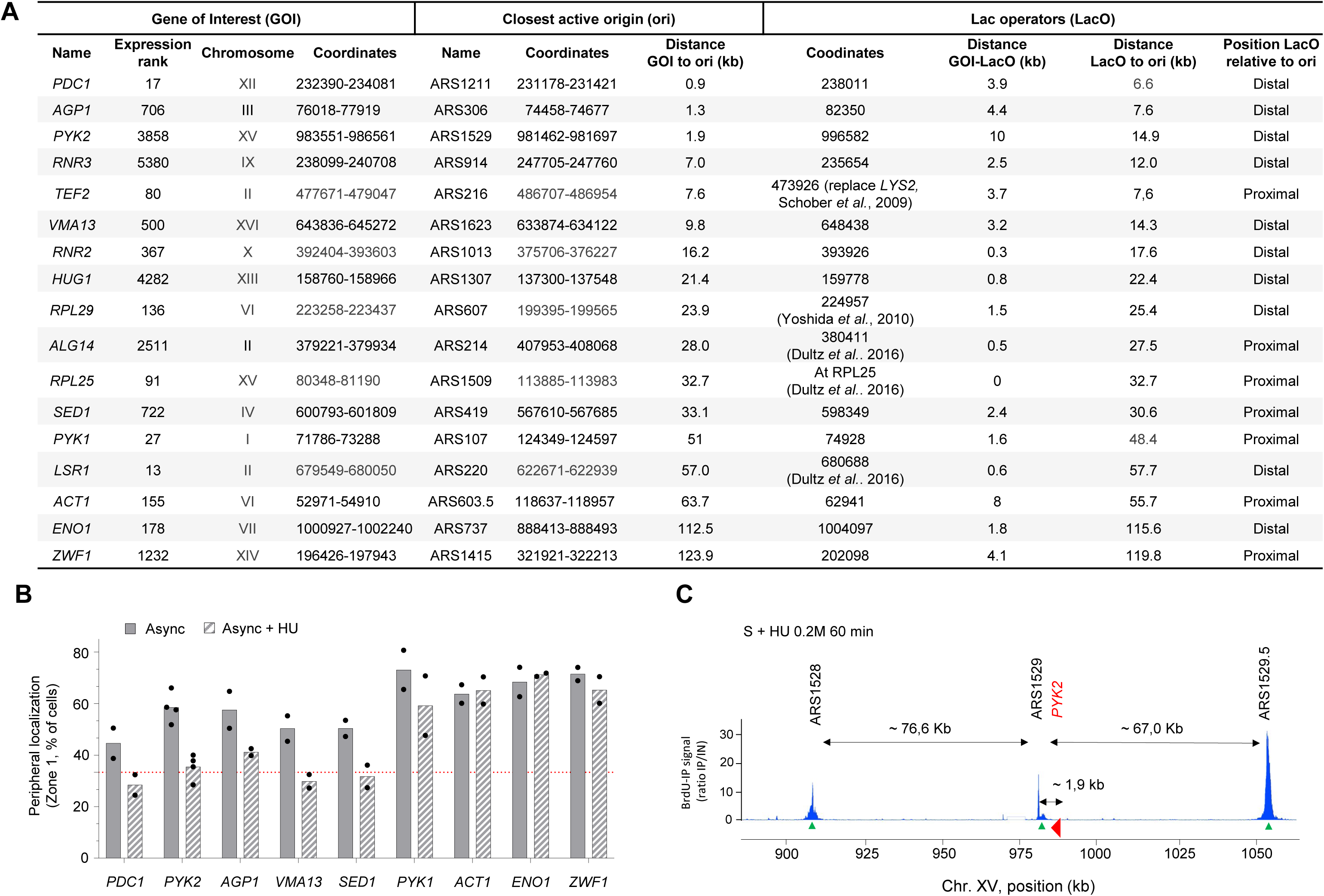
Gene release from nuclear pores is coupled to DNA replication progression. (A) Summary of genes of interest and *lacO* insertion sites. (B)Percentage of GFP foci in Zone 1 under the indicated conditions (Async = exponentially growing cells, Async + HU = exponentially growing cells treated with 0.2 M HU for 60 min). Independent biological replicates are indicated by data points (N≥3, >100 cells/biological replicate). The horizontal dashed red line indicates 33% (random distribution). Statistical comparisons were performed using pairwise Fisher’s exact test (*P < .05, **P < .01, ***P < .001, ****P < .0001). (C) Illustration of the replication profile (BrdU-IP) on chromosome XV in wild-type cells after α-factor synchronization and release into 0.2 M HU for 60min. Positions of early-firing origins are indicated by green arrowheads. The *PYK2* locus is depicted in red. Approximate distances are indicated.

**Figure S3:**
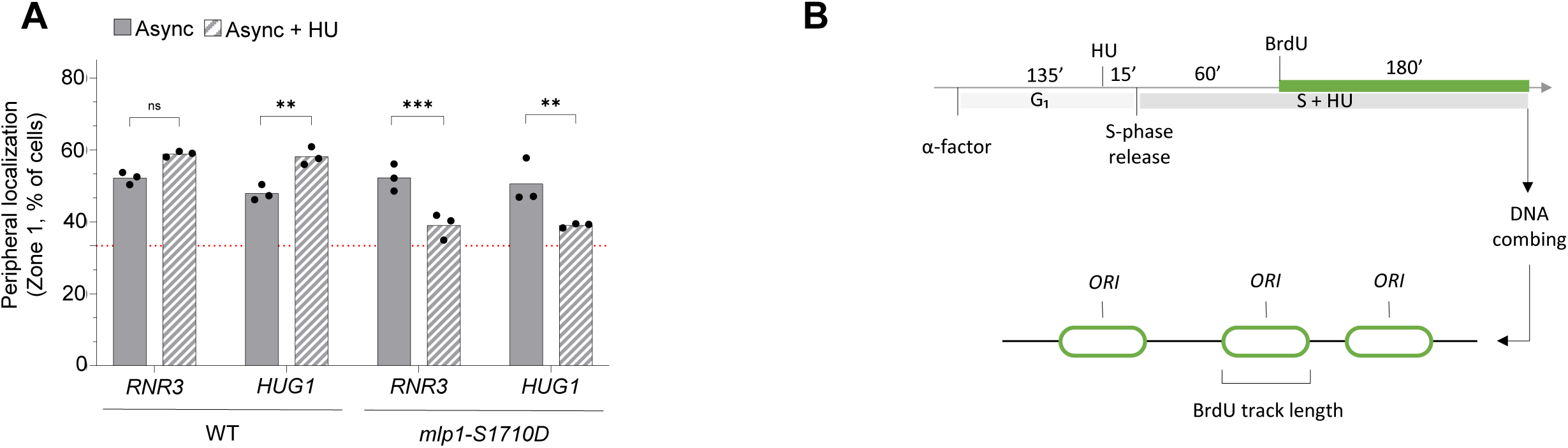
*mlp1-S1710D* prevents reassociation of genes to NPCs during replication stress. (A) Percentage of GFP foci in Zone 1 for *RNR3* and *HUG1* in the indicated strains in exponentially growing cells treated with 0.2 M hydroxyurea for 60 min (Async + HU) or left untreated (Async). Independent biological replicates are indicated by data points (N≥3, >100 cells/biological replicate). The horizontal dashed red line indicates 33% (random distribution). Statistical comparisons were performed using pairwise Fisher’s exact test (*P < .05, **P < .01, ***P < .001, ****P < .0001).

## Experimental procedures

### Yeast strains, culture conditions and flow cytometry

All strains used are listed in Table S1. For liquid cultures, synthetic complete medium was supplemented with 2% glucose unless otherwise stated. *MAT***a** cells were synchronized in G1 by adding α-factor (5 μg ml^-1^, GeneScript) for 120 min at 25°C unless otherwise stated. Arrest without buds was monitored by phase microscopy. G1-blocked cells were released into S phase by the addition of pronase (75 µg ml^-1^, Millipore) and were treated or left untreated with 0.2 M HU (Angene, 127-07-1). The SLD3-SLD7-CDC45 inducible construct was induced by addition of β-estradiol (1µg µl^-1^, Sigma-Aldrich) 1 h prior to release into S phase. Rapamycin (1µg ml^-1^, Interchim) was added 30 min prior to cell harvest in experiments using the anchor-away technique.

### Drop assays

Exponentially growing cells were counted using a CASY cell counter (OLS system) and were diluted or concentrated to 1.10^7^ cells/ml. 10-fold serial dilutions were spotted onto YPAD or SC plate +/-the indicated drugs. Where indicated 5-fold dilution series were used. Plates were incubated for 2 to 7 days at 25°C or 30°C.

### Protein extracts and western blotting

TCA-precipitated protein extracts were resolved by SDS-PAGE using a custom-made low cross-linking gel for the detection of the Rad53 mobility shift, and transferred to a PVDF membrane using a Bio-Rad Trans-Blot Turbo Transfer system (30 min, standard mode). After blocking, proteins were probed with a rabbit polyoclonal anti-Rad53-phospho (dilution 1/3000; a gift from C. Santocanale, University of Galway). Following washing, membranes were incubated with HRP-conjugated anti-rabbit IgG secondary antibody (dilution 1/5000; Cytiva NA934-1ML). Blots were scanned with a ChemiDoc MP (BioRad) imaging system and semi-quantitative determination of protein level was performed using the Image J (Fiji) software.

### DNA combing

*S. cerevisiae* cells expressing codon-optimized TK and ENT1 were grown to 6 x 10^6^ cells/ml in YPD medium, synchronized in G1 and released into S-phase with 0.2 M HU. Cells were labeled with 400 µg/mL BrdU (B5002, Sigma) added 75 minutes post-release. Cells were harvested in 0.1% sodium azide, washed, and treated with Zymolyase buffer (50 mM phosphate buffer pH 7, 50 mM EDTA pH 8, 1.2 M Sorbitol, 10 mM DTT, 2.5 mg/mL Zymolyase 20T, 07663-91, Nacalai Tesque). Cells were embedded in 2% low-melting agarose (LMA, 11AGAL0050, MPBiomedicals) plugs (90 µL/plug), digested in Proteinase K buffer (125 mM EDTA pH 9.5, 1% Sarkosyl, 1 mg/mL Proteinase K) at 37 °C for 24 hours, and extensively washed. Plugs were equilibrated in MES buffer (1X MES pH 6, 100 mM NaCl) and melted at 67 °C. Genomic DNA was digested overnight with β-agarase (M0392L, NEB) at 42 °C. DNA was combed onto silanized coverslips (10490802, PolyAn) using an automated pulling machine (250 µm/s, Genomic Vision), baked at 60 °C for 2 hours, and mounted on glass slides. Coverslips were denatured with 1 M NaOH for 20-22 minutes, blocked, and incubated sequentially with primary antibodies against BrdU (347580, Becton Dickinson, 1:20) followed by fluorescent secondary antibody Alexa 546 (A-11030, Sigma, 1:50), and ssDNA (1:10) followed by fluorescent secondary antibody Alexa 647 (A-21241, Invitrogen, 1:50), with intermediate washes. Slides were mounted in Prolong Gold (P36930, Invitrogen). Images were acquired using a Zeiss Axio Imager Z3 microscope equipped with a Hamamatsu ORCA-Flash 4.0 LT+ camera. Track lengths were measured using MetaMorph software, and data were plotted using Python, with 100 to 200 tracks analyzed per sample. To evaluate reproducibility and variability, SuperPlots were generated. Statistical significance was determined using a one-way ANOVA on replicate median values followed by Tukey’s multiple-comparisons test (ns, p > 0.05; *p < 0.05; **p < 0.001; ***p < 0.0001) across three biological replicates, as described by Lord et al. (2021).

### Microscopy and image analysis

Cells were fixed in fresh 4% (w/v) paraformaldehyde (PFA) for 1 min, washed 3 times in PBS and attached to a #1.5H (170 µm) glass coverslip using concanavalin A. They were imaged by acquiring 50 z-slices of 200 nm thickness. A Zeiss Axio imager Z2 was used to acquire images, equipped with a Hamamatsu ORCA-Flash 4.0 LT+ camera, a Märzhäuser XY motorized stage, a Prior NanoScanZ piezo, and a Plan apochromat 100x/1.46 oil objective. GFP and tdTomato were excited using a XCite 120 LED, Zen-driven LED light source. Fluorescence images were deconvolved using Huygens Professional software using the Classic Maximum Likelihood Estimation (CMLE) algorithm with a signal-to-noise ratio of 1, automatic background estimation and 40 iterations. Quantification of foci in nuclear zones was done using the PointPicker plug-in (http://bigwww.epfl.ch/thevenaz/pointpicker/) on Fiji by measuring the position of the DNA locus spot in respect to the nuclear periphery. The nucleus is divided in 3 concentric zones of equal volume and the DNA locus is sorted into one of the zones based on the relative distance of the DNA locus spot to the nuclear periphery (Da mota *et al*., 2024). At least 300 cells/condition were scored (>100 cells/biological replicate).

## Data availability

No deposited datasets are included.

## Acknowledgements

We thank the Montpellier Ressources Imagerie (MRI) imaging facility. We thank Elisa Dultz, Karsten Weis, Susan Gasser, and Jason Brickner for sharing zoning strains and Amir Aharoni for sharing plasmids to insert *lacO*. G.N. thanks the FNR for a PhD fellowship (project ID 18862584). J.H thanks the Ligue Contre le Cancer for a PhD fellowship. This work was supported by the Agence Nationale de la Recherche (ANR-22-CE12-0031-01, “SCooTR”) and IUF grants (Junior, 2022) to J.P.

## Author contributions

Conceptualization: GN, JH and JP. Investigation: GN, JH, SG, LF, AB and JP. Methodology: GN, JH and JP. Formal analysis: GN and JP. Validation: GN and JP. Visualization: GN and JP. Funding acquisition, Supervision and Project administration: JP. Writing – original draft – GN and JP. Writing – review & editing: GN, JH, PP and JP.

## Declaration of interests

The authors declare no competing interests.

## Notes

### Competing Interest Statement

The authors have declared no competing interest.

